# Phenotypic heterogeneity in a batch culture of *Chlamydomonas reinhardtii* with different light tolerances

**DOI:** 10.1101/2025.07.30.667606

**Authors:** Gaganpreet K. Gill, Dion G. Durnford

## Abstract

Genetic diversity of populations is essential for generating phenotypic variation to allow a flexible response to a shift in environmental conditions. Therefore, in populations of genetically identical individuals grown in the lab, you would predict that phenotypic heterogeneity would be small. However, we isolated two subpopulations of genetically identical individuals from an exponentially growing batch culture of the microalga *Chlamydomonas reinhardtii* using Percoll step-gradients. The culture fractionated into a low-density, Top fraction and a high-density, Bottom fraction. These subpopulations displayed several phenotypic differences, including size, protein content, the amount of chlorophyll per cell, and photosynthetic performance. Because of the variation in pigment content and photosynthetic performance, we tested the hypothesis that there are differences in their tolerance to light stress. Following high-light stress, the Bottom subpopulation was more resistant to photodamage, had a greater capacity for light dissipation, and had a minimal photoacclimation response to high light, compared to the Top subpopulation. The Bottom population also had a greater resistance to exogenously induced singlet oxygen stress mediated by rose bengal. We hypothesize that these subpopulations are derived from stochastic mechanism where the Bottom subpopulation has activated a general high-light stress response pathway as part of a “bet-hedging” strategy that could give it a fitness advantage with a shift towards a light-stress environment.

## Introduction

Genetic and phenotypic diversity are crucial for microbial populations to acclimate to their environments and adapt to new ecological niches. Natural populations coexist within communities that are shaped by biotic interactions between different species and abiotic factors, such as chemical or physical properties of the environment that change seasonally or unpredictably (Kassen and Rainey, 2004; Ackermann, 2015). Environmental heterogeneity is expected to drive genetic and species diversity in populations (Kassen, 2002; Kassen and Rainey, 2004). Unique niches in these heterogeneous environments induce selective pressures, whereby advantageous traits may become more prevalent, driving survival, adaptation, and evolution of phenotypically diverse populations living in naturally dynamic environments (Kassen and Rainey, 2004).

Unexpectedly, phenotypic variation can occur within isogenic populations that are grown under the same environmental conditions. This phenotypic heterogeneity has been observed in a variety of microorganisms. Variation in individuals can come about via molecular noise, where small variations in regulatory molecules can have effects on the phenotype (Korobkova et al., 2004), or where epigenetic regulation of regulatory proteins create phenotypic variation, as observed with yeast adhesion proteins (Halme et al., 2004). Phenotypic variation can also stem from differences in cell age as cells undergo asymmetric separation of cell components during cell division that can produce phenotypic differences. In *Escherichia coli* (*E. coli*), the asymmetric distribution of pole proteins during division can give rise to physiologically younger cells with newly synthesized poles or older cells with the original pole composition (Stewart et al., 2005). The “older” cells have a slower growth rate and reduced resilience compared to the “younger” cells.

Phenotypic heterogeneity is also observed in yeast where oxidatively damaged proteins are unevenly distributed between mother and daughter cells during binary fission in synchronized cells (Erjavec et al., 2008; Roux et al., 2010). The unequal distribution of damaged proteins ultimately “sacrifices” one daughter cell so the other can thrive (Barker and Walmsley, 1999; Aguilaniu et al., 2003; Lynch, 2005; Fredriksson and Nyström, 2006). Daughter cells inheriting damaged proteins aged more rapidly, had longer generation times, and had reduced longevity as compared to their “younger” siblings (Erjavec et al., 2008; Roux et al., 2010). In yeast, asymmetrical distribution can also include the concentrations of metabolites, signalling pathway components (Colman-Lerner et al., 2005; Woldringh, 2005; Rang et al., 2011; Wang et al., 2022), and damaged cellular components, such as peroxisomes (Choudhry et al., 2018), that can lead to cell-to-cell heterogeneity in isogenic populations. This has the effect of creating individuals in a population with different functional capacities, and a heterogeneous population from genetically identical individuals.

The nature to which this functional heterogeneity exists in other model eukaryotic systems, such as microalgae, is unclear, especially with respect to the functional significance. For instance, with the microalga *Chlamydomonas reinhardtii* (*C. reinhardtii*), isogenic batch cultures are commonly used in experiments, and despite these cultures starting from a single individual, *Chlamydomonas* cultures have variation in traits such as lipid content, cell size, starch metabolism, gene expression, and growth rates (Garz et al., 2012; Lee et al., 2013; Damodaran et al., 2015;). Lee et al. (2013) observed stochastic heterogeneity in lipid content by encapsulating single *Chlamydomonas* cells in alginate hydrogel microcapsules and stained them with the lipid-binding fluorescence stain, BODIPY, revealing significant variation. Similarly, Garz et al. (2012) used second harmonic generation signals and laser scanning microscopy to quantify starch content per cell in synchronized, isogenic cultures and found that metabolic processes related to starch synthesis and degradation are major contributors to the heterogeneity. These findings suggest that intrinsic factors can drive phenotypic diversity in isogenic cultures of *Chlamydomonas*.

Heterogeneity in *Chlamydomonas* has also been observed as variations in growth dynamics. Despite using isogenic *Chlamydomonas* batch cultures in consistent conditions, Damodaran et al. (2015) found significant growth-rate heterogeneity using a millifluidic drop analyzer (a device that tracks the number of cells over time) to assess the growth kinetics. Using this device, the authors identified two subpopulations - the major subpopulation was termed as, “fast-growing” cells, and the minor subpopulation as “slow-growing” with doubling times of 7-11 hours and 12-17 hours, respectively. Damodaran et al. (2015) suggests this may be due to some isogenic cells expressing an active dividing state, while others express a restricted dividing state. These studies collectively show that *Chlamydomonas* populations, like bacterial and yeast cultures, develop heterogeneity and phenotypic diversity due to unknown, intrinsic mechanisms.

Building on the understanding of phenotypic heterogeneity in microorganisms, we wanted to investigate this phenomenon in an isogenic batch culture of *C. reinhardtii* cells with the goal of isolating different subpopulations, determine how they phenotypically differ, and determine whether the phenotypic variation was functionally significant. We attempted to physically isolate *Chlamydomonas* subpopulations using Percoll density gradient centrifugation, a technique used to separate microalgal cells based on buoyant density (Whitelam et al., 1983; Lavoie et al., 1986). Here we describe the separation of two populations that have differences in macromolecular composition, growth, photosynthesis and respiration. Importantly, these subpopulations showed differences in their tolerance to high light, suggesting they may have a different fitness under changing environmental conditions.

## Materials and Methods

### Growing and synchronizing batch cultures

*C. reinhardtii* strain CC125 was obtained from the *Chlamydomonas* Resource Center at the University of Minnesota (https://www.chlamycollection.org/). Cultures were maintained on Tris-Acetate Phosphate (TAP; <u>Harris, 1989)</u> agar plates under continuous light at room temperature until ready for use.

To ensure an isogenic isolate, we streaked the culture on a TAP plate and isolated a single colony for further propagation. Liquid starting cultures were initiated by aseptically inoculating cells from the TAP agar plate into a 125 ml Erlenmeyer flask containing 35 ml of sterile TAP liquid media. This starting culture was closed with a foam stopper and wrapped with aluminum foil and grown for four days with constant shaking in a 13-hour light and 11-hour dark (13/11-hour L/D) cycle (Hlavová et al., 2016). The light intensity of the growth chamber was 75 µmol quanta m^-2^s^-1^ using cool-white, fluorescent bulbs, and the cultures were grown on a shaker at 24°C inside a growth chamber. The starter culture was then used to inoculate new cultures in fresh TAP media at a starting cell abundance of 3 x 10^5^ cells ml^-1^. These experimental cultures were returned to the same growing conditions for two days. Unless otherwise indicated, experimental cultures were harvested four hours into the light period (L4) two days following culture dilution.

### Percoll density gradient centrifugation

Density gradient centrifugation using a Percoll step gradient (Sigma-Aldrich, Saint Louis, U.S.A.) was used to separate subpopulations of cells from cultures. The gradients were made by overlaying 2 ml of 40% Percoll™ (Cytiva, U.K.), 6.5 ml of 25% Percoll, and 3 ml of 5% Percoll in a 13.2 ml open-top, ultra-clear ultracentrifuge tube (Beckman Coulter, Inc, Fullerton, CA, U.S.A.). Percoll solutions were diluted with water. Gradients were loaded with a total of 2.5 x 10^7^ cells in a 0.5 ml volume and centrifuged at 7885 x g for 30 minutes at 24°C using a SW41 rotor in an Optima XE-90 Ultracentrifuge (Beckman Coulter, Inc, Fullerton, CA, U.S.A.). Subpopulations (or “bands”) were collected using a syringe and needle and kept in sterile falcon tubes.

The subpopulations were washed with media from the original culture (spent media) to remove any residual Percoll and cells collected by centrifugation at 8050 x g for 15 minutes (15°C). Pelleted cells were then resuspended in known volumes of spent media and used for different measurements.

### Cell abundance, Chlorophyll, and starch quantification

A sample of cells were fixed with Lugol’s iodine (Throndsen, 1978) and counted using a Neubauer hemocytometer under a light microscope.

Chlorophyll was extracted from 2 x 10^6^ cells in 1 ml of 100% methanol and pigments quantified by measuring absorbances at 470, 652, 665, and 750 nm using a Beckman Coulter DU720 UV-Vis spectrophotometer as previously described by Zamzam et al. (2022).

For starch analysis, the starch pellets were thawed, suspended in 200 µL of 80% ethanol, and boiled for 20 minutes to solubilize the pellet. Starch content was determined using a Total Starch Assay Kit (AA/AMG Megazyme, Ireland) according to the manufacturer’s instructions and as described by Zamzam et al. (2022).

### Protein extraction and quantification

Samples of each subpopulation were collected by centrifugation (15,000 x g for 10 minutes) and the pellets containing whole-cells were stored at −80°C until ready for analysis. Protein extraction and quantification have been previously described (Meagher et al, 2021).

### Confocal fluorescence microscopy

A qualitative assessment of lipid droplets in the subpopulations was done using nile red (Sigma) and confocal microscopy. Samples of each subpopulation (1 x 10^6^ cells ml^-1^) were fixed with 10 µL of 4% paraformaldehyde (PFA) and stained with nile red (Sigma) to a final concentration of 1 µg/ml (in methanol) and incubated in the dark for one hour. The images were captured using a Leica TCS SP8 confocal LSM (analyzed using LAS-X software) and processed using ImageJ (Fiji). Lipid droplets were visualized using an excitation laser 552 nm, and emission at 500-610 nm. Chlorophyll autofluorescence was simultaneously visualized using an excitation laser 488 nm, and emission at 650-710 nm.

### Chlorophyll fluorescence and Photosynthesis Measurements

Photosynthetic efficiency was evaluated by measuring chlorophyll fluorescence using a pulse amplitude modulated PAM101 fluorometer (Walz, Germany). Sample preparation and measurements were done as described by Damoo and Durnford (2021).

Photosynthetic activity was evaluated by measuring oxygen evolution rates using an oxygen electrode and Oxygraph (Hansatech Instruments Ltd, U.K.). The oxygen evolution rates were calculated using O2View software (Hansatech Instruments Ltd) and the data was normalized to cell number, then plotted as a Photosynthesis-Irradiance (PI) curve.

### Flow cytometry

#### Cell size

A S3e Cell Sorter (Bio-Rad Canada) was used to measure the cell sizes of subpopulations. Forward scatter (FSC) and side scatter (SSC) measurements were taken of a minimum 1 x 10^5^ cells of each subpopulation. The average cell size of the subpopulations was estimated using a standard calibrated microsphere bead set (Spherotech Inc., USA) as described by Zamzam et al. (2022). Using FlowJo^TM^ Software, an initial gate was created by selecting the population of interest on the linear FSC-Area (A) and SSC-A density plot (supplementary Fig. S1, panel A). From the gated population, doublets were excluded by further gating the dense population of events along either a linear FSC-Height (H) and FSC-W density plot, or a linear FSC-A and FSC-H density plot (supplementary Fig. S1, panel B). The average modes for unique FSC-W scale values were used to generate a standard curve (supplementary Fig. S2), and the subpopulation’s cell diameters were calculated using the linear regression line.

### DNA content

DNA content was analyzed by staining cells with a double-stranded DNA-binding fluorescent stain, Vybrant^TM^ DyeCycle^TM^ Green stain (5 mM in DMSO) (Thermo Fisher Scientific). This stain is excited by a 488 nm laser with a 534/34 band pass filter. Unstained samples were used as a control to account for background fluorescence.

### Growth rate

Relative growth rates over the first two days of transfer to fresh media were determined using absorbance at 750 nm (A750) in cultures grown in 12-well microwell plates.

Subpopulations were diluted with sterile fresh media to a final concentration of 3 x 10^5^ cells ml^-1^ and two ml aliquots were placed into individual wells of the microwell plates. Samples were grown for three days (13/11-hour L/D cycles, 24°C) on a shaker and the absorbance was measured at 750 nm using a SpectraMax® M3 Multi-Mode microplate reader (Molecular Devices, U.S.A.) at least twice a day as a proxy for cell abundance. Yield rates were calculated using modified expressions from Liu (2017):

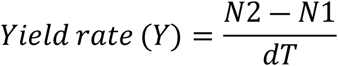

For a qualitative assessment of growth yield, the subpopulations were diluted to a starting concentration of 4 x 10^6^ cells ml^-1^ and 10 µl of each sample was pipetted onto a solid TAP media plate and grown for four days to look for visual cues for differences in their growth.

### Rose bengal treatment

Rose bengal (RB) is a photosensitizer that induces singlet oxygen formation when exposed to light (Ledford et al., 2007). Samples of each subpopulation (200 µl of 1 x 10^6^ cells ml^-1^ suspensions) were added to 96-well microplates with 40 µl of increasing concentrations of RB. The plates were incubated either in the dark (control) or under light (75 µmol quanta m^-2^s^-1^) for 1.5 hours, then cells were collected and centrifuged for 10 minutes at 15,000 x g (13°C). The supernatants containing RB were removed and the pelleted cells were resuspended in 240 µl of fresh TAP media. For a qualitative assessment, 5 µl of the samples were pipetted onto a solid TAP media plate and grown for up to 4 days under continuous light (56 µmol quanta m^-2^s^-1^).

### Statistical analyses

Data was imported into Microsoft Excel^TM^ for graphing and computing Student *t*-tests (two-tailed, two-sample equal variance). Two-way ANOVAs and Tukey’s HSD tests were performed using RStudio (RStudio Team, 2023).

## Results

### Two subpopulations separated based on buoyant density

We explored heterogeneity of isogenic *Chlamydomonas* batch cultures by fractionating cells based on buoyant density. The cultures were harvested at Day 2, four hours into the light period (L4). These cells are growing exponentially, thus avoiding any significant age-related differences in the culture.

When we centrifuged the culture on a Percoll step gradient (5-25-40%) it consistently fractionated into two subpopulations: a lower density population (Top) at the interface of the 5-25% Percoll boundary; and a higher density population (Bottom) at the 40%-Percoll step. The Top subpopulation was significantly more abundant with 86% ± 7.9% SD of the cells compared to the Bottom subpopulation (with 14% of the cells; p=1.6 x 10^-27^, Student’s *t*-test) (Fig. 1).

**Figure 1.**
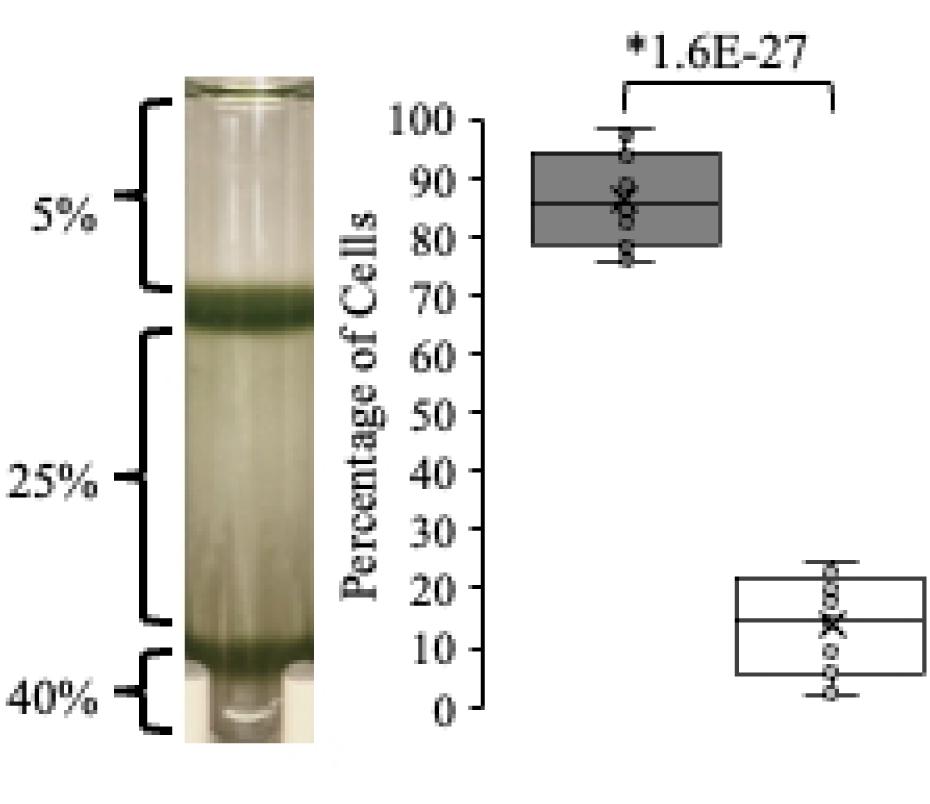
Separation of a batch culture into two subpopulations using a Percoll step-gradient. Proportions of the Top (gray box) and Bottom (white box) subpopulations are shown as box and whisker plots with the median shown as the center line. Individual data points (circles, n=19) are shown. Asterisks indicate significant differences between the bands with the p-value indicated (Student’s /-test).

To determine the stability of the fractions, the bands were collected, washed, and re-centrifuged onto new Percoll step-gradients (Fig. 2A). In this test, cells from the Top subpopulation again fractionated at the 5-25% Percoll interface and the Bottom subpopulation at the 25-40% interface. This confirmed a robust separation based on density and helped confirm no additional subpopulations diverged from either band (Salvo et al., 1982; Maurer et al., 2022).

**Figure 2.**
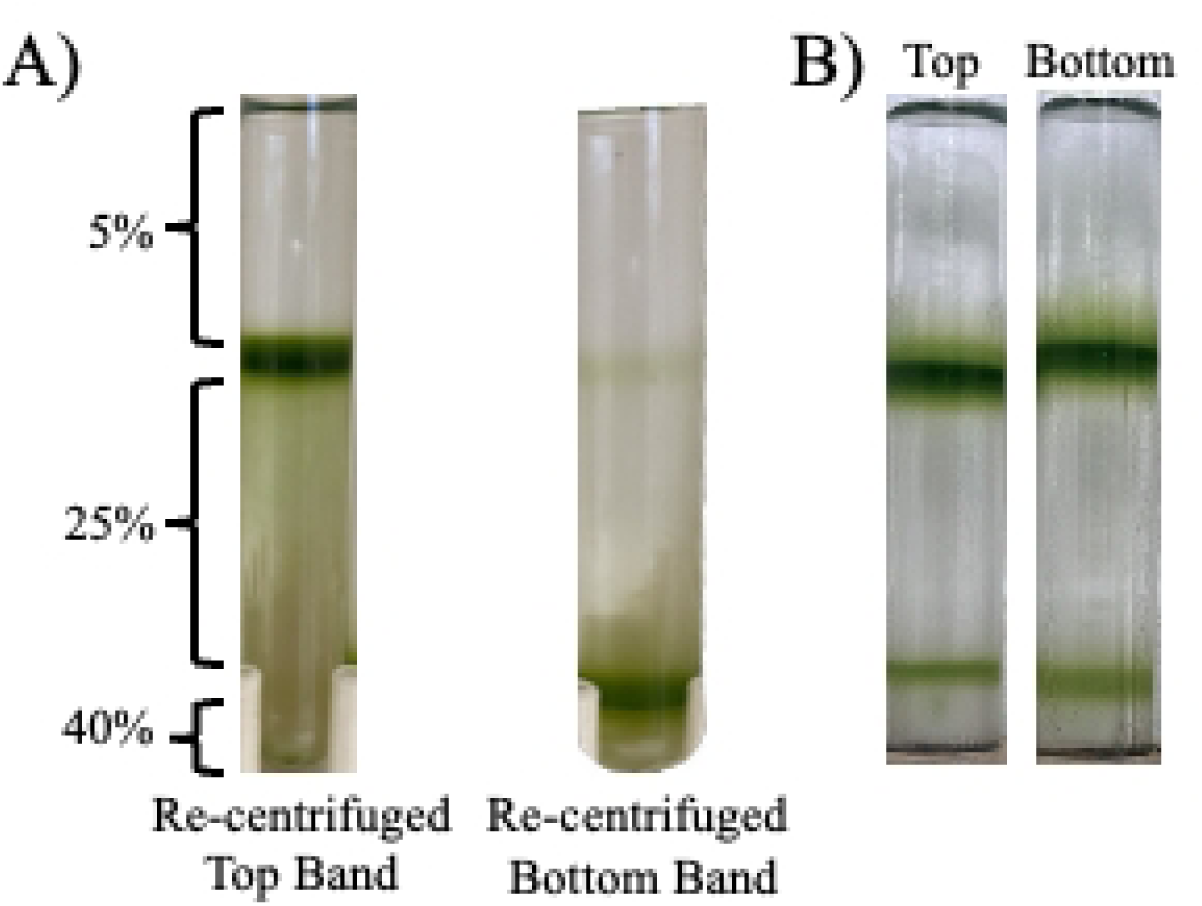
Stability of the subpopulations A) Each subpopulation collected and recentrifuged onto new Percoll step-gradients, and B) Top and Bottom subpopulations were collected and inoculated into fresh TAP media liquid cultures (35 ml) starting at 5.4 x 10^4^ cells ml^-1^. Subpopulations were grown for three days and loaded onto new Percoll step-gradients.

Although we started this study by isolating a single colony to ensure the CC125 culture was isogenic, we tested the possibility that the Top and Bottom subpopulations may be fixed phenotypes in the culture, whether due to genetic variation or epigenetic modification (Fraser and Kærn, 2009; Ackermann, 2015). Each subpopulation was collected and inoculated into new flasks with TAP media and grown for three days under the same 13/11-hour L/D regime. In each case, a Top and Bottom fraction were identified in the same, approximate relative distribution of both subcultures (Fig. 2B).

To test if the subpopulations represented different stages of the cell cycle we first monitored the Day 2 culture over a 24 hr period of the L/D cycle to see if the culture was synchronized. Other than an increase from the start of the light period to hour 2, there was no significant increase in cell abundance during the light period (Fig. 3). At the start of the dark period, however, cell abundance increased about 1.5-fold indicating about half of the cells divided. The cells did accumulate chlorophyll over the light period, suggesting cells were getting larger, and the chlorophyll per cell declined when cells started to divide in the dark phase (Fig. 3). Overall, this indicated that the culture was reasonably synchronized. To specifically test whether the Top and bottom fractions differed in DNA content, we stained each subpopulation with Vybrant^TM^ DyeCycle^TM^ Green stain to assess DNA content per cell using flow cytometry. We did not observe a difference in DNA content between the Top and Bottom fractions that would suggest variation in stage of the cell cycle or explain differences in buoyant density (Fig. 4).

**Figure 3.**
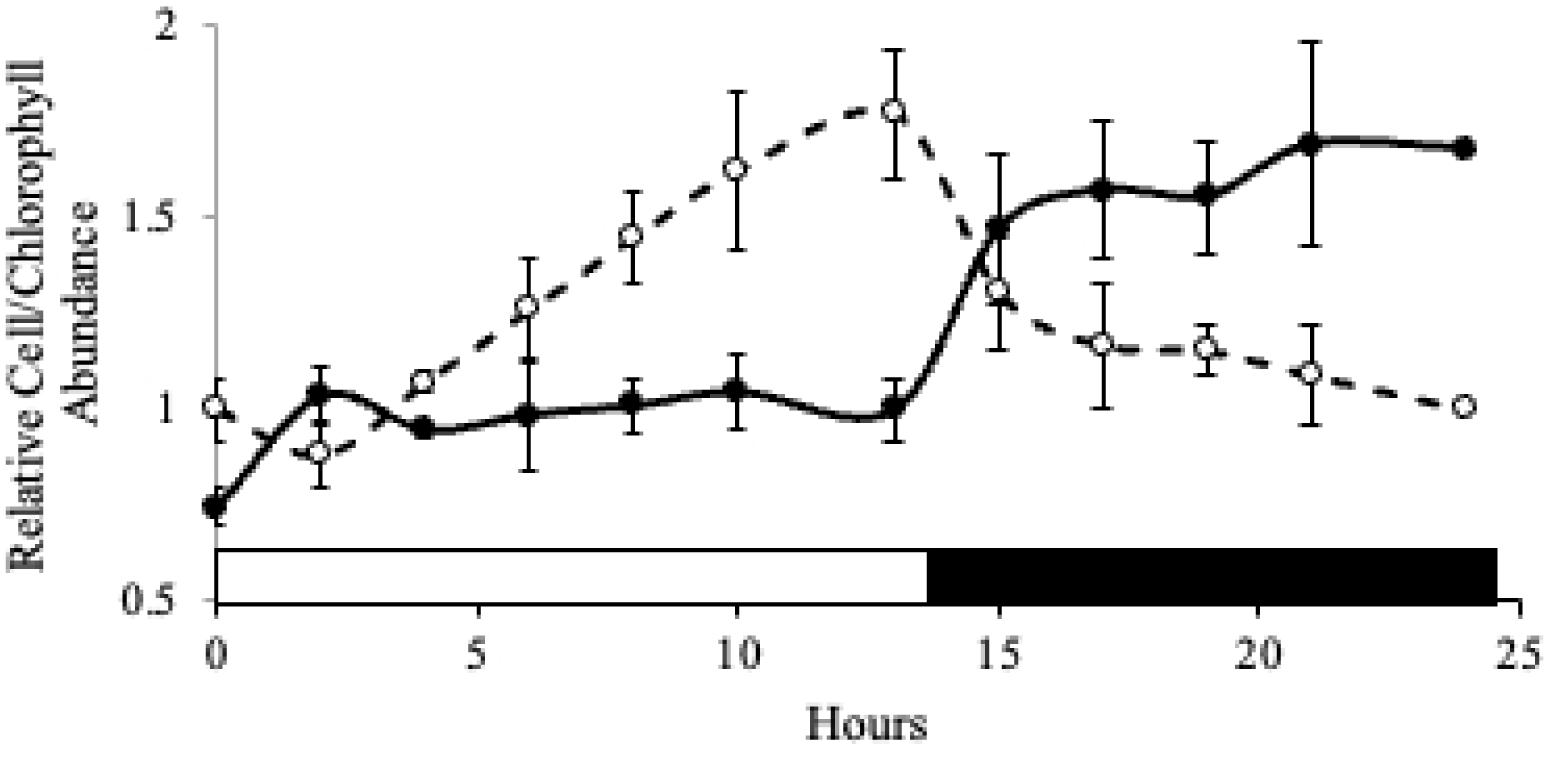
Relative cell abundance and chlorophyll content per cell over one L’D cycle in a Day 2 batch culture. Relative number of CC125 cells per ml (closed circles, solid line; n=3, +SD for all except L13 (n=6) and Dll (n=2)). Relative amount of total chlorophyll content *(a+b,* pg cell-^1^) (open circles, broken line). The open bar on the bottom of the graph indicates light period, and the closed bar indicates the dark.

**Figure 4.**
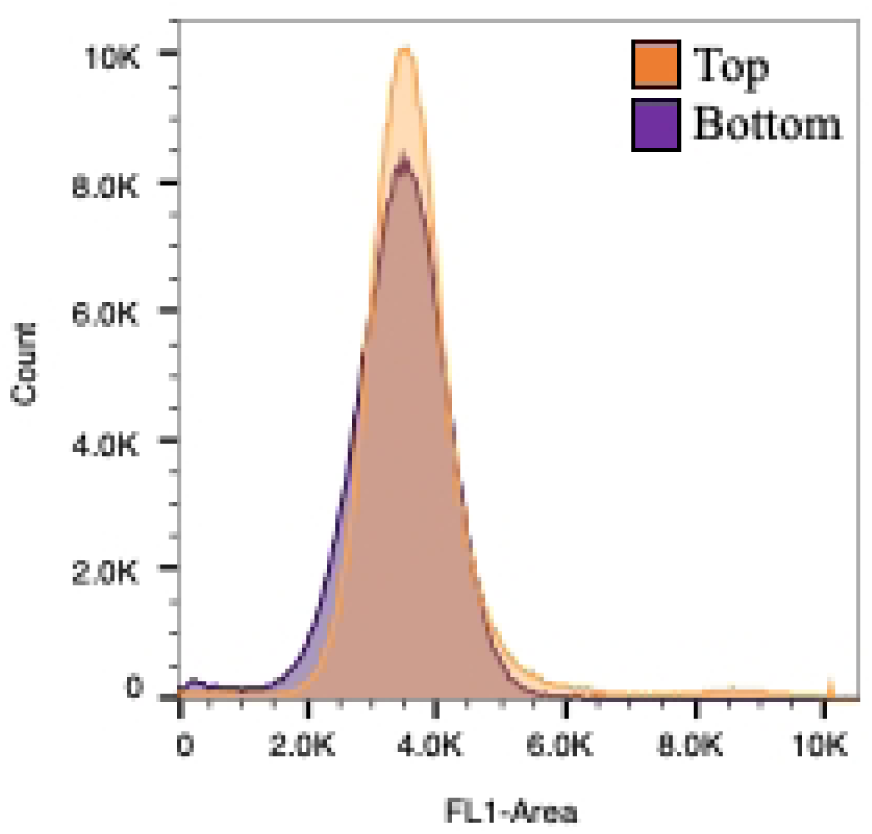
DNA fluorescence of the Top and Bottom subpopulations using flow cytometry that were harvested at L4 on Day 2. Subpopulations were stained with Vybrant DyeCycle Green and the histograms of DNA Fluorescence-Area for both subpopulations are overlayed.

To decipher the differences between the Top and Bottom subpopulations, we initially focused on morphological characteristics, such as cell size and shape. Using brightfield microscopy, we surveyed 300 cells from each fraction and observed 89% of the cells in the Bottom fraction were predominantly ellipsoidal, compared to 68% in the Top fraction. While there was clearly a variation in morphology in each subpopulation, the cells in Figure 5 were chosen from a dozen confocal microscopy images and represent the cell size and shape most observed. The chlorophyll autofluorescence panels show the typical arrangement of a cup-shaped chloroplast along the bottom of the cells, and the ellipsoidal-cells had smaller plastid lobes. In Figure 4, nile red was used to stain lipid droplets in the cells and its fluorescence was examined using confocal microscopy. The rounder cells in the Top fraction had more lipid droplets compared to the more elliptical cells in the Bottom fraction.

**Figure 5.**
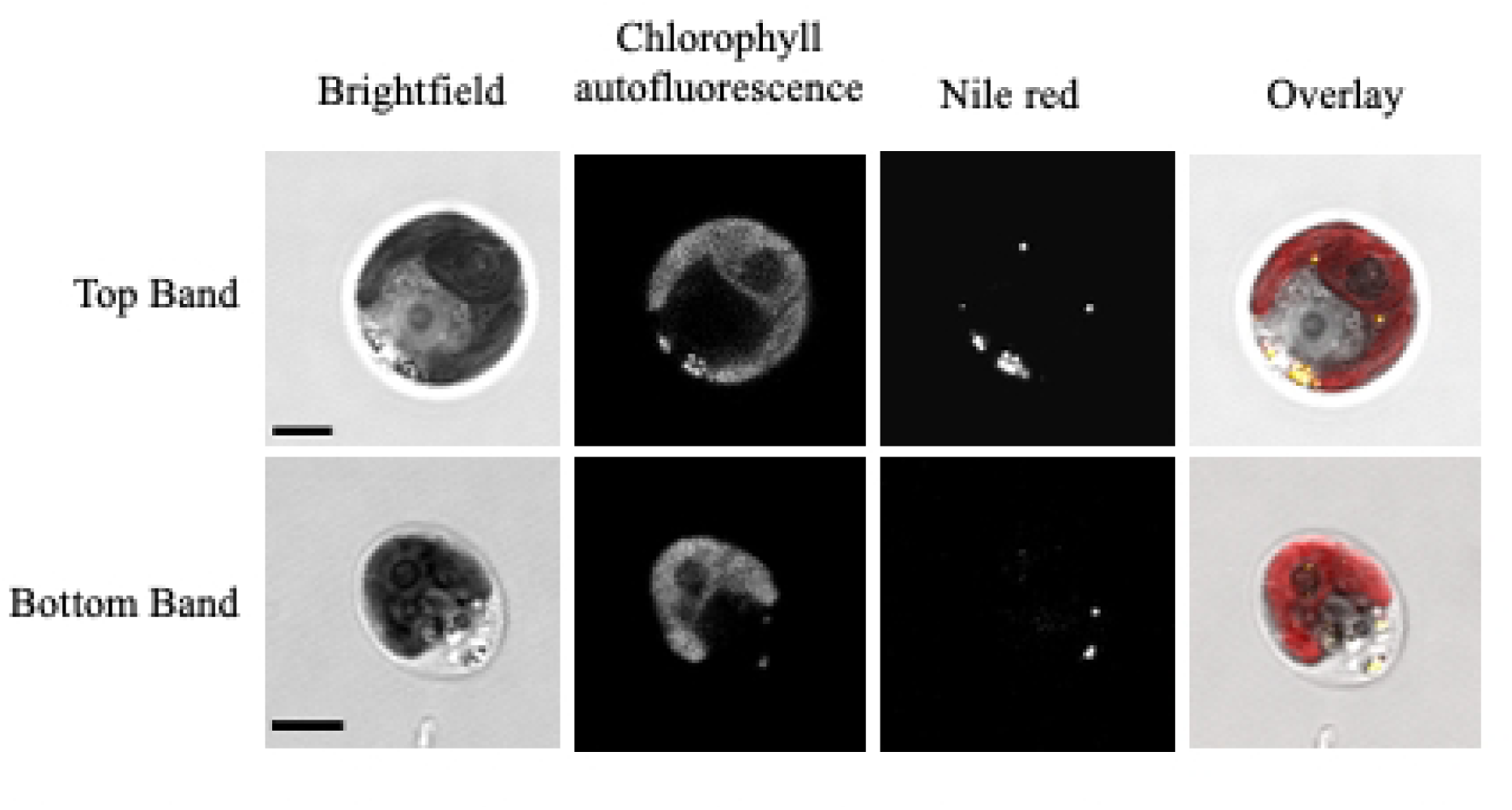
Morphology of the most observed cell types in each subpopulation. Chlorophyll autofluorescence microscopy shows the shape of the plastid and Nile Red-staining of each subpopulation detects lipid droplets using fluorescence microscopy. For the overlay images, nile red (yellow) and chlorophyll autofluorescence (red) are shown. Scale bar 5 µm. Images were enhanced for brightness and contrast using ImageJ.

The different populations had clear differences in morphological and biochemical properties. Using flow cytometry, we measured the cell diameter (Table 1), and on average the Top subpopulation was significantly larger than the Bottom population. Assuming the volume of a sphere, this equates to volume difference of 14% between the two populations. The total protein content per cell was about 40% greater in the Top subpopulation than the Bottom (Table 1). The total chlorophyll content per cell was similarly higher in the Top subpopulation on a per cell basis, though the ratio of Chl *a* and *b* in the two subpopulations was identical (Table 1). Despite the differences in protein and chlorophyll content, there were no differences in the amount of starch per cell between the two subpopulations (Table 1).

**Table 1.** Size and biochemical characteristics of the Top and Bottom subpopulations.

| Characteristic | Top | Bottom |
| --- | --- | --- |
| Cell diameter ($\mu\text{m}$) | $9.1 \pm 0.4$ * | $8.7 \pm 0.3$ |
| Total chlorophyll *a+b* ($\text{pg cell}^{-1}$) | $6.0 \pm 1.1$ * | $4.0 \pm 1.3$ |
| Chl *a* ($\text{pg cell}^{-1}$) | $4.4 \pm 0.8$ * | $2.9 \pm 0.9$ |
| Chl *b* ($\text{pg cell}^{-1}$) | $1.6 \pm 0.4$ * | $1.0 \pm 0.3$ |
| Chl *a/b* ratio | $2.9 \pm 0.3$ | $2.9 \pm 0.3$ |
| Total protein ($\text{pg cell}^{-1}$) | $171 \pm 15.6$ * | $110.1 \pm 19.7$ |
| Total starch ($\text{pg cell}^{-1}$) | $2.5 \pm 1.3$ | $1.6 \pm 1.0$ |

\*significantly different from Bottom subpopulation ( $p < 0.006$ , Student's *t*-test). Error bars are ± SD ( $n=12$ for cell diameter; $n=19$ for chlorophyll content; $n=5$ for protein and starch content)

### Functional characteristics of the subpopulations

#### Growth rate

To determine if there were differences in growth rate between the Top and Bottom subpopulations, we conducted quantitative growth assay on media and a qualitative assay on agar plates. The Top subpopulation had an 18% greater average yield rate (number of cells added to the population over time) over two days, where the Top population grew by 8.7 x 10^4^ cells hr^-1^ (± 2.2 x 10^4^ SD) while the Bottom population grew by 7.1 x 10^4^ cells hr^-1^ (± 2.1 x 10^4^ SD, p=0.01, Student’s *t*-test) (Fig. 6A). When equal numbers of cells were plated onto a TAP plate and observed after four days, the growth differences are visible and in agreement with the liquid assay (Fig. 6B).

**Figure 6.**
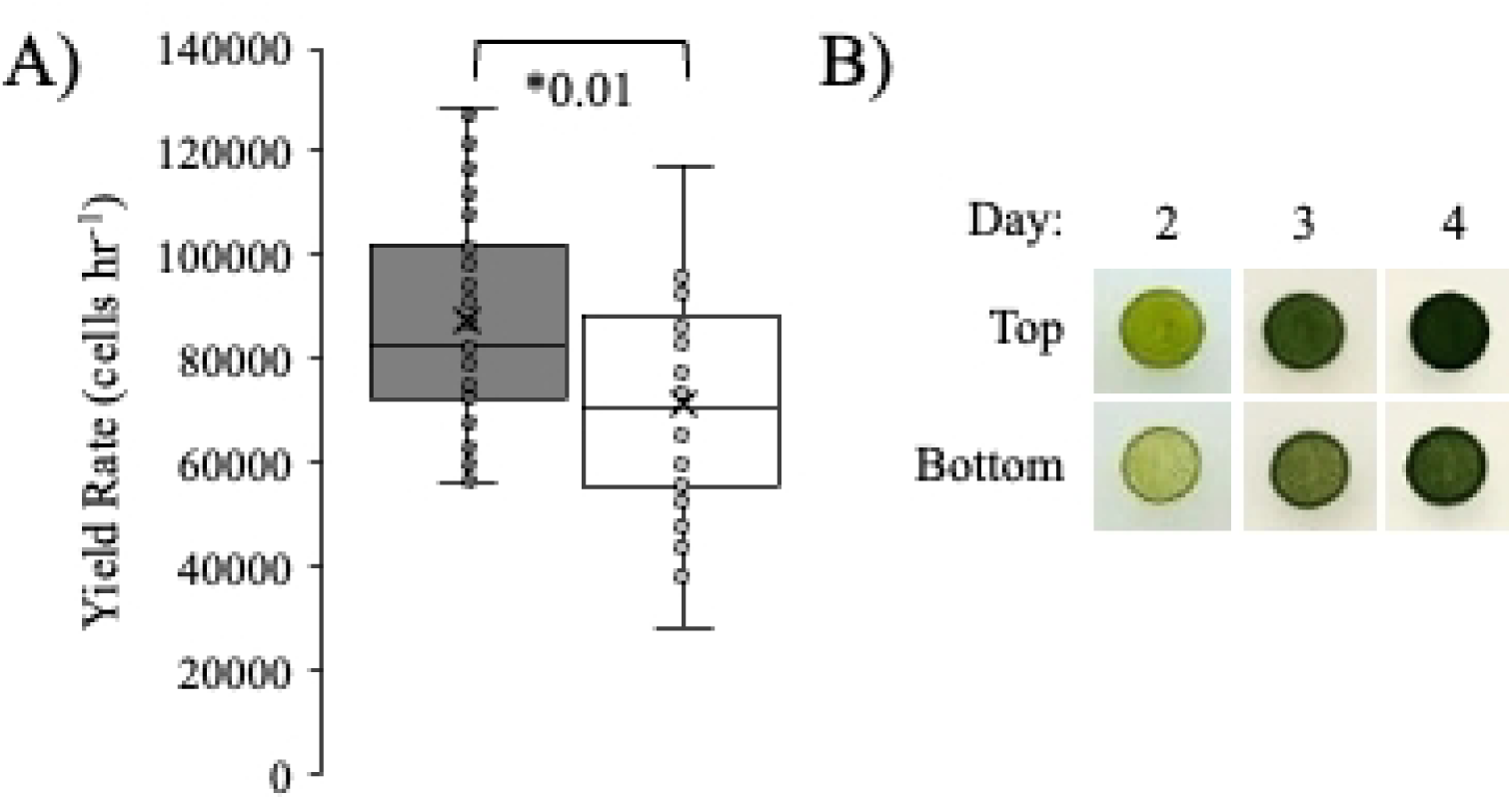
Growth differences between the Top and Bottom subpopulations. A) Average yield rates after growth in liquid media for 2 days. The boxes represent 50% of the data and the whiskers show the distribution of the upper and lower 25%. Asterisks indicate significant differences between the bands with the p-value indicated (Student’s /-test). Individual data points (Top band, filled circles, n=26) (Bottom band, open circles, n=25) are shown. B) Subpopulations grown on TAP agar plates for 2-4 days as a qualitative assessment of growth rate.

### Photosynthesis

There were also differences in photosynthetic performance between the subpopulations and the Top subpopulation had a significantly greater maximum photosynthetic rate (Pmax) on a per cell basis (Table 2; p=0.001, Student’s *t*-test) and a greater light-use efficiency (α) (p=0.002, Student’s *t*-test), which was determined by the initial slope of the PI curve (Walker, 1987). The dark oxygen consumption rate, which is a proxy for mitochondrial respiration, was almost double the amount in the Top subpopulation than in the Bottom subpopulation (p=0.05, Student’s *t*-test), suggesting a greater metabolic activity in the Top fraction, explaining the faster yield rate.

**Table 2.**
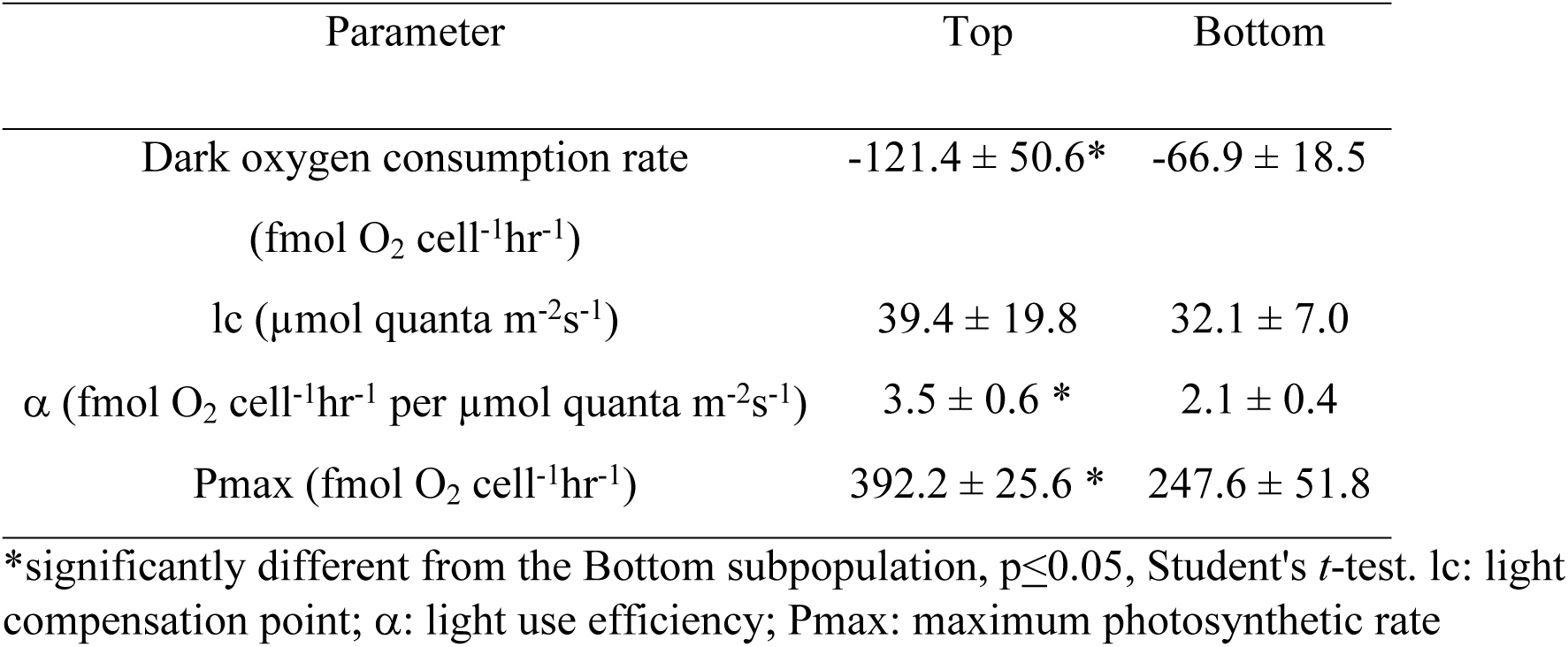
Photosynthetic parameters of the Top and Bottom subpopulations (n=5, ±SD).

### High-light stress tolerance

Considering the differences in photosynthetic rates and chlorophyll levels, we examined the photoprotection and light dissipation capacity using chlorophyll fluorescence (Fig. 7). We first tested the light tolerance of each subpopulation in a short-term light stress experiment. Cells from each subpopulation were diluted to 2-5 x 10^6^ cells in 10 ml of spent media and placed in 50 ml Erlenmeyer flasks. Subpopulations were incubated in low-light (LL) (75 µmol photons m^-2^s^-1^) or high-light (HL) (530 µmol photons m^-2^s^-1^) for 2 hours with constant shaking, after which the Fv/Fm was determined (Fig. 7A). There was no difference in the Fv/Fm between the populations under LL, but there was a small, significant difference in the Fv/Fm after HL-exposure. The Fv/Fm of the Top subpopulation was 0.17 ± 0.05 SD, while the Fv/Fm of the Bottom fraction was 0.25 ± 0.05 SD (p=0.01, Student’s *t*-test). This suggested that the Top subpopulation was more photoinhibited during the HL exposure.

**Figure 7.**
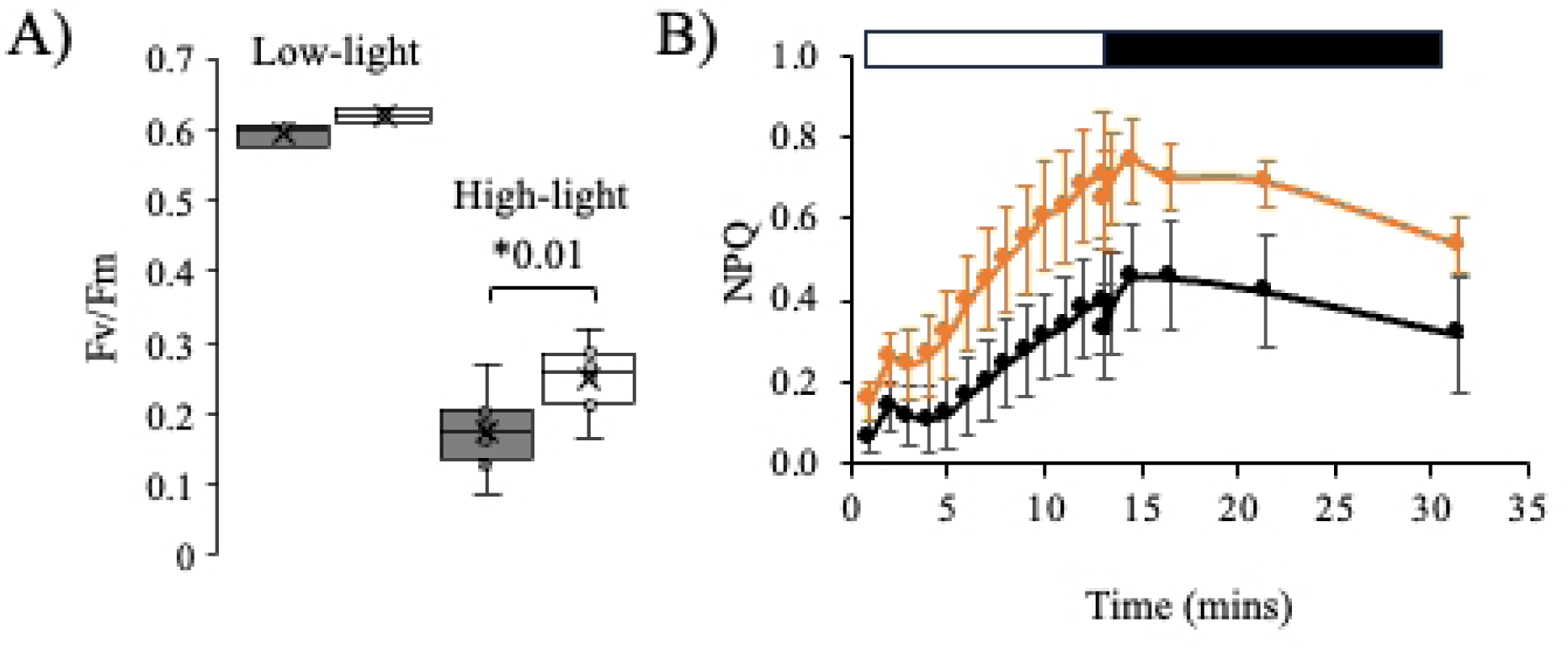
Photosynthetic efficiency and maximum quantum efficiency of PSII in the Top and Bottom subpopulations. A) Subpopulations were exposed to LL (75 µmol quanta m^-2^s^-1^) or HL (530 µmol quanta m^-2^s^-1^) for two hours and all contents of the flasks were measured for their Fv/Fm. The boxes represent 50% of the data and the whiskers the distribution of the upper and lower 25%. Asterisks indicate significant differences between the Top (gray) and Bottom (white) subpopulations with the p-value indicated (Student’s /-test). B) Average light dissipated by non-photochemical quenching (NPQ) over time for Top (black) and Bottom (orange) subpopulations (n 6, ± 2SE). The open bar indicates the hours spent in light, and the closed bar indicates hours spent in the dark.

Since there seemed to be differences in tolerance to HL-stress, we assessed the capacity of each population to dissipate light energy by measuring non-photochemical quenching (NPQ) during an equivalent HL-stress. NPQ is a way to bypass light energy away from the core reaction centre to protect the photosynthetic apparatus (Demmig-Adams et al., 2014). In these experiments, the Bottom subpopulation showed a significantly greater amount of light dissipated through NPQ than the Top subpopulation (Fig. 7B). At 10 minutes, the average amount of light dissipated through NPQ in the Bottom subpopulation was double (0.605 ± 0.2 SD) that of the Top subpopulation (0.314 ± 0.1 SD, p=0.006, Student’s *t*-test) (Fig. 7B). During the dark recovery period, both populations showed a similar recovery with evidence for photoinhibition at the end of the recovery period (Fig. 7B).

Next, we examined each subpopulation’s capacity to photoacclimate to HL stress over the course of a 13-hour period using chlorophyll content as a marker for photoacclimation state (Fig. 8A). Typically, *Chlamydomonas* cells increase in size over the course of the day (Hlavová et al., 2016; Heldt et al., 2020). As the cell gets bigger, the amount of chlorophyll content per cell also increases (as seen in Fig. 3) and we can follow that as a marker of light responses.

**Figure 8.**
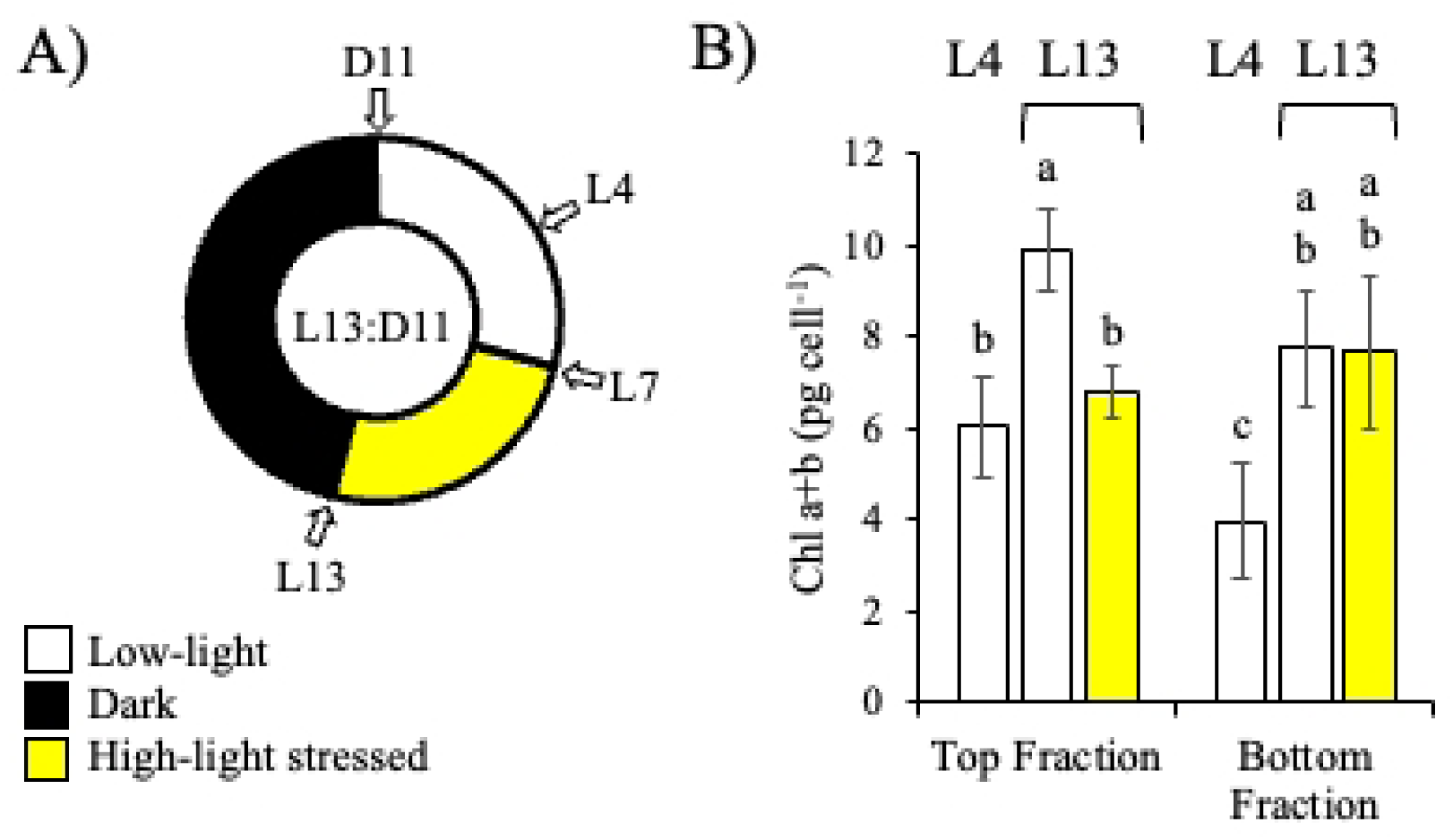
Effects of high-light stress on Top and Bottom subpopulations. A) Diagram illustrating the experimental design with or without a HL (531 µmol quanta m^-2^s^-2^) exposure of the batch culture between L7-L13 on Day 2. At LI 3, the culture was fractionated on a Percoll gradient to separate out the Top and Bottom fractions. B) Total chlorophyll content *(a+b)* for each subpopulation was measured at photoperiods L4 and L13 (n 3-6, ±SD) (n=19 for L4). Bars labelled with different letters indicate significant differences (p<7e-5 for ANOVA, and p<0.05 for Tukey’s HSD test).

With a light shock experiment (Fig. 8A), the Top and Bottom populations responded differently to excess light. Under four hours of LL (L4), the Top fraction had a greater chlorophyll content per cell than the bottom (Fig. 8B), as previously noted (Table 1). Continuing under LL conditions, both fractions increased their chlorophyll content over the course of the light period by 69 and 96% for the Top and Bottom subpopulations, respectively. Although, the Top fraction still had the higher chlorophyll content per cell at the end of the day (L13; 22% greater, Fig. 8B). When a HL shock was administered starting seven hours into the light period (L7) and lasting until the end of the day (L13), chlorophyll accumulation in the Top population was significantly repressed (6.8 pg cell^-1^ ± 0.5 SD; p<0.05, Tukey’s HSD test) (Fig. 8B), while the HL-exposed Bottom fraction’s chlorophyll content remained unchanged compared to its LL control (Fig. 8B). There was no significant difference in the total number of cells present for either subpopulation under LL or HL (supplementary Fig. S3).

### Oxidative stress tolerance

We also tested each subpopulation’s ability to tolerate oxidative stress to help decipher differences in the photoprotection pathways seen thus far. We used increasing concentrations of RB, a photosensitizer that induces the formation of singlet oxygen when it is exposed to the light (Gorman and Rodgers, 1992; Ledford et al., 2007; Miao et al., 2019). In the light, RB ultimately transitions from its ground state (S0) to an excited triplet state (T1) (Gorman and Rodgers, 1992). In the triplet state, RB transfers its energy to ground-state oxygen which excites molecular oxygen into its singlet state (^1^O_2_*). In *Chlamydomonas*, accumulation of singlet oxygen is toxic and can induce photodamage. In this experiment, cells were exposed to increasing concentrations of RB for 1.5 hours in the light or the dark (control), to simulate a progressively stringent singlet oxygen stress. After which, cells were washed and plated onto a fresh TAP plate to screen qualitatively for survival. In the dark, both the Top and Bottom subpopulations were unaffected, as expected (Fig. 9). However, in the presence of light the Top subpopulation was considerably more sensitive to the RB-treatment, with obvious effects starting at 14 µM. This indicates that the Bottom subpopulation was less sensitive to singlet oxygen stress.

**Figure 9.**
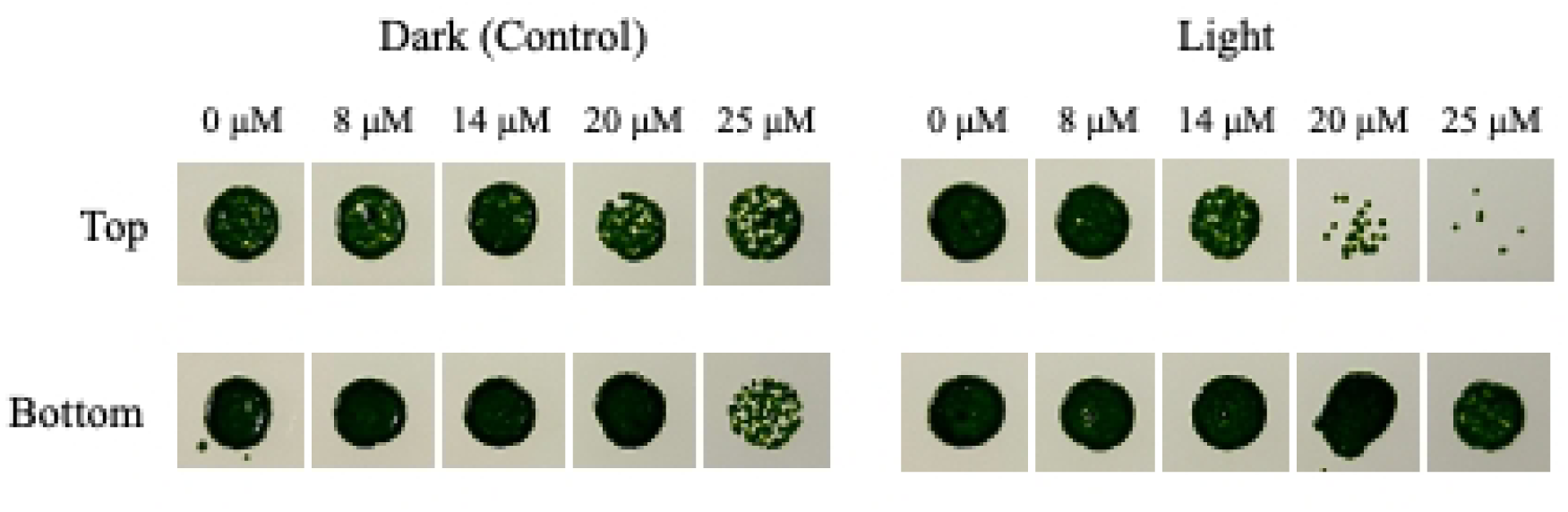
Rose Bengal (singlet oxygen) resistance of the Top and Bottom subpopulations. Subpopulations were diluted to a starting concentration of 1 x 10^6^ cells ml^4^, exposed to rose bengal in the presence or absence (control) of light and 5 µl of the culture was dispensed onto TAP agar plates. Subpopulations grown on TAP agar plates for 7 days as a qualitative assessment of growth after exposure to singlet oxygen from rose bengal treatment.

## Discussion

### Heterogeneity in natural and isogenic microbial populations

Microbial populations in nature will exhibit genetic and phenotypic diversity even when individual cells share the same habitat. But diversity still arises because ecosystems are naturally dynamic, undergoing changes like temperature fluctuations, nutrient depletion, seasonal changes, and shifts in resource availability. To survive, organisms must adapt to these environmental fluctuations which can result in phenotypic diversity amongst cells of the same species. On the other hand, when isogenic cells are grown under controlled laboratory conditions, they are expected to show minimal phenotypic variation because the uniform environment should elicit cells to behave identically across the whole population. However, in our study, we consistently detected at least two subpopulations in batch cultures of *C. reinhardtii* (CC125) using Percoll density gradient centrifugation, which separates cells based on buoyant density.

Previous studies have shown evidence of cellular heterogeneity in isogenic cultures of *Chlamydomonas* through biochemical or physiological characteristics such as cell size, starch content (Garz et al., 2012), lipid content (Lee et al., 2013), and growth rates (Damodaran et al., 2015). Similarly, in isogenic bacterial populations, heterogeneity has been identified through asymmetric cell division that causes unequal inheritance of cellular components, such as old cell poles (Stewart et al., 2005; Woldringh, 2005), or the accumulation of damaged intracellular proteins (Fredriksson and Nyström, 2006). The spontaneous formation of persister cells in bacteria are another example of heterogeneity arising from isogenic populations, where a portion of the population switches to a dormant phenotype that is resistant against antibiotics (Balaban et al., 2004; Kussell et al., 2005; Smith et al., 2018). In isogenic yeast populations, heterogeneity has been observed as variations in replicative aging (Barker and Walmsley, 1999), the accumulation of carbonylated proteins (Erjavec et al., 2008; Roux et al., 2010), cell cycle arrest in older cells, and differences in signaling pathways (Colman-Lerner et al., 2005; Wang et al., 2022). These findings demonstrate how phenotypic variation and cellular heterogeneity are not restricted to natural environments but can also manifest in isogenic populations under controlled conditions, consistent with the results observed in this study.

### Buoyant density differences of Top and Bottom subpopulations are independent of cell cycle stage

It wasn’t clear what was responsible for the differences in buoyant density between the two populations. Buoyant density is a tightly regulated cellular property essential for maintaining key biological processes including division, differentiation, and apoptosis (Yancey et al., 1982). Cells of the same cell-type typically maintain a consistent density emphasizing its functional importance (Kubitschek et al., 1983; Kubitschek, 1987; Osypiw et al., 1994; Allen et al., 2006; Bryan et al., 2010; Neurohr and Amon, 2020; Kispert et al., 2024). The observed differences in buoyant density between the Top and Bottom subpopulations may arise from several factors, including variations in water content, osmoregulation (Baldwin and Kubitschek, 1984; Kubitschek, 1987; Baldwin et al., 1988; Kültz, 2001), organelle composition (Kubitschek, 1987; Bazzani et al., 2021), and metabolic state (Yancey et al., 1982; Hare et al., 1998; Jogawat, 2019; Ghosh et al., 2021). In *Chlamydomonas*, osmoregulation is also mediated by a contractile vacuole (CV) complex that has two CVs in the anterior-end, each of which is formed by the fusion of numerous small vesicles before fluid expulsion (Gruber and Rosario, 1979; Luykx et al., 1997). Since CV size is linked to cell size in *Chlamydomonas*, with larger cells having longer intervals between systole contractions and a greater pumping rate of fluid expulsion (Luykx et al., 1997; Komsic-Buchmann et al., 2014), the larger Top subpopulation may have higher water content contributing to its lighter buoyant density, but this is something we cannot confirm. Structural differences, such as variations in cell wall or chloroplast composition (Bazzani et al., 2021) may also contribute to differences in density between subpopulations.

Additionally, osmolytes are solutes like carbohydrates, polyols, and amino acids, are critical for cellular osmoregulation and stress adaptation (Yancey et al., 1982; Hare et al., 1998; Jogawat, 2019; Ghosh et al., 2021). In *Chlamydomonas*, osmolytes such as glycerol and proline accumulate in response to environmental stresses like HL (Duval et al., 1999; Davis et al., 2013) and salinity (Reynoso and De Gamboa, 1982; Mastrobuoni et al., 2012; Park et al., 2020). If the Bottom subpopulation in our study is predisposed of greater levels of osmolytes, like proline, it may contribute to their greater buoyant density and enhanced resistance to HL stress as compared to the Top subpopulation.

While buoyant density can change as a culture ages, as seen in *E. coli* cultures (Makinoshima et al., 2002), we avoided this complication by consistently harvesting cells on the same day post inoculation and at the same point in the photoperiod (L4). We also ruled out differences in the cell cycle being a major factor in the buoyant density, as can be observed in some yeast cultures (Bryan et al., 2010), as there were no differences between the two subpopulations in DNA content per cell at the period we isolated cells. There was also no evidence of genetic differences that would explain the variation in buoyant density, which would be expected to be a more permanent feature of the population. First, we isolated the strain from a single colony to ensure it was isogenic. Second, we also looked at the stability of the Top and Bottom phenotype by initiating cultures directly with each fraction. This ultimately led to the similar distribution of the fractions in the population after three days (Fig. 2B). This finding aligns with Damodaran et al. (2015) who showed inoculating cultures using their fast-growing *Chlamydomonas* subpopulation produced heterogenous progeny, rather than exclusively-fast growing cells. While the reason for the differences in buoyant density are not clear, the evidence points to the idea that cells are consistently responding to culture conditions, intrinsic factors, or some sort of internal signal that is triggering the phenotypic differences (Garz et al., 2012).

### Functional differences

The different subpopulations have different capacities for light use, which was reflected in the differences in chlorophyll content and photosynthetic parameters. The Top subpopulation has a greater light use efficiency and greater maximal photosynthetic rate. This, coupled with a greater dark respiration, should provide growth advantages especially under light-limiting conditions, and in general, we found a faster yield rate in short-term growth experiments. The Bottom fraction, however, was better able to dissipate excess light, effectively being in a more high-light acclimated state, which agrees with the lower chlorophyll content in the cell, a useful indicator of light acclimation status (Bonente et al., 2012; Humby et al., 2013).

The Bottom subpopulation had phenotypic characteristics that should give it an adaptive advantage under HL stress. This was confirmed in short-term (2 hours) HL-stress experiments where the Bottom subpopulation showed a greater ability to resist photoinhibition estimated by looking at the maximum quantum efficiency of PSII after the HL-shock. The Bottom subpopulation also was better able to dissipate excess light energy through non-photochemical quenching (NPQ), a way to de-excite chlorophyll and redirect energy away from the rection centre to avoid photodamage (Demmig-Adams et al., 2014).

The Bottom subpopulation also showed more resistance in response to high light over a six-hour time frame when applied mid-light phase. At the end of the light period (L13) the Bottom subpopulation exposed to HL continued to accumulate chlorophyll, as did the LL control. But, cells in the Top fraction showed a decline in chlorophyll content per cell, showing a typical response to HL exposure (Webb and Melis, 1995; Shapira et al., 1997; MacIntyre et al., 2002; Durnford et al., 2003; Tokutsu et al., 2021). However, the mechanism of this chlorophyll reduction in HL can be complex. While it is likely due to an inhibition of the synthesis and accumulation of chlorophyll-binding proteins in response to HL, it could also be triggered by increasing cell division where the chlorophyll is diluted as the cells divide. This wouldn’t be surprising given that *Chlamydomonas* diverts excess light energy into metabolic pathways that can act as energy sinks (Falkowski and LaRoche, 1991; Davis et al., 2013) and could trigger cell division. As more cells reach the critical size (Craigie and Cavalier-smith, 1982; Heldt et al., 2020) and undergo cell division by L13. However, we were unable to resolve any significant differences in cell number following HL exposure. Of course, with the Bottom subpopulation, we didn’t see the decline in chlorophyll, which would imply that compositional and functional differences in the cells, including an enhanced NPQ capacity, minimized light stress such that chlorophyll accumulation was not inhibited over the light period. We also did not observe any differences in cell abundance between the LL and HL samples in the bottom subpopulation, which indicates that cell division was not triggered. Nevertheless, there was a clear difference in the response of the two subpopulations to HL exposure, signifying significant functional differences in their response to light.

In addition to a more muted response to HL stress, the Bottom subpopulation also had an increased tolerance to a singlet oxygen stress induced by rose bengal (Fig. 9). When *Chlamydomonas* is exposed to too much light beyond what it can dissipate, there is the inevitable production of ROSs that can damage biomolecules and become toxic to the cell (Erickson et al., 2015). Cells have a variety of inducible detoxification and repair mechanisms as part of an ROS acclimation response (Erickson et al., 2015; Ghosh et al., 2021). For example, Ledford et al. (2007) found enhanced resistance to singlet oxygen in *Chlamydomonas* when cells were exposed to RB under HL, suggesting an overlap between ROS and HL-induced response pathways. The acclimation response is partly regulated by the SAK1 gene in *Chlamydomonas*, which becomes phosphorylated in the presence of singlet oxygen (Wakao et al., 2014; Erickson et al., 2015). The greater tolerance of RB in the bottom population may mean a more global tolerance of stressors related to excess light, and perhaps the pathways related to ROS detoxification (Triantaphylidès and Havaux, 2009), or at least those for singlet oxygen. The enhanced singlet oxygen resistance could also be related to other mechanisms, such as the accumulation of osmolytes or other chemicals that act as ROS scavengers as part of a global HL-stress response mechanism (Alia et al., 1991, 2001; Hare and Cress, 1997). Overall, cells in the Bottom subpopulation have attributes resembling a HL-acclimated state (Falkowski and LaRoche, 1991; Nawrocki et al., 2020; Dupuis et al., 2025), making them more resistant to HL and singlet oxygen damage.

### Origin of phenotypic heterogeneity

The source of the phenotypic heterogeneity in a batch culture of genetically identical individuals grown under the same conditions is intriguing. Phenotypic variation could arise from cellular aging which contributes to heterogeneity over time (Stewart et al., 2005; Woldringh, 2005; Choudhry et al., 2018) or through the accumulation of damaged molecules (Erjavec et al., 2008; Roux et al., 2010) that mimic aging processes by reducing the fitness and viability of cells over time (Książek, 2010). While we did use young, two-day old cultures and standardized the harvesting time to make these processes less likely, it’s difficult to rule out such mechanisms. Currently, mechanisms contributing to any cell division asymmetry in *Chlamydomonas*, whether age or stress related, are unknown.

Environmental factors, both abiotic and biotic, can contribute to phenotypic variation in isogenic populations (Rouag and Dominy, 1994; Smith et al., 2018; Saccardo et al., 2022), but in our experiments we attempted to keep conditions in the flask uniform. However, it is difficult to rule out distinct microenvironments in our cultures that could activate specific signal pathways in subsets of the population (Kærn et al., 2005; Blake et al., 2006; Fraser and Kærn, 2009; Ackermann, 2015; Davis and Isberg, 2016; Gasperotti et al., 2020), though we would have predicted more variability between experiments if that were the case (Rouag and Dominy, 1994; Saccardo et al., 2022).

Cell-to-cell or neighbor interactions could also drive phenotypic switching. For example, in bacteria autoinducers (signal molecules) can trigger phenotypic-switching response based on the cell abundance. This may, for instance, cause a change in biofilm formation (Barak and Ulitzur, 1981; Yarwood et al., 2004; Hense and Schuster, 2015) or secretion of virulence factors (proteins) (Dunman et al., 2001) and increase the variability in the population. However, such systems are not known in *Chlamydomonas,* to our knowledge.

Finally, stochastic mechanisms, including random gene expression and metabolic fluctuations, play a key role in generating phenotypic heterogeneity (Gasperotti et al., 2020). Single-cell transcriptome studies in *Chlamydomonas* have found significant variability in the expression of photosynthesis-related, stress-response, and metabolic genes (Garz et al., 2012; Damodaran et al., 2015; Bheda, 2020). This variability can be driven by epigenetic regulation or stochastic phenotypic switching (Lee et al., 2013; Bheda, 2020), which allows individual cells to toggle between distinct gene expression states (Hung et al., 2014), potentially explaining the observed differences in the photoprotective strategies and ROS tolerance between the Top and Bottom subpopulations. For example, the ROS detoxification mechanisms and acclimation responses in the Bottom subpopulation may be linked to stochastic gene regulation of commonly regulated stress-response pathways, that may involve of SAK1 (Wakao et al., 2014), for instance. Further investigation using techniques such as RNAseq and/or single-cell sequencing on the Top and Bottom populations could provide deeper insight into the stochastic mechanisms and underlying phenotypic variation in *Chlamydomonas* batch cultures.

In isogenic cultures of the microalga, *Haematococcus pluvialis*, cells can stochastically switch between mobile and non-mobile phenotypes with different stress tolerances (Tang et al., 2024). Random fluctuations in gene expression that present as gene noise can cause phenotypically identical cells to differ in behaviour as well, due to varied protein or enzyme levels, which in turn affects metabolic pathways and other functions (Elowitz et al., 2002; Levin, 2003; Korobkova et al., 2004; Kilfoil et al., 2009; Pinheiro et al., 2022; de Groot et al., 2023).

Although isogenic cells experience a baseline level of genetic noise, being a biological system, they still present as phenotypically identically cells. When this noise surpasses a certain threshold, it can lead to phenotypically distinct subpopulations that are now detectable based on their traits (Elowitz et al., 2002; Blake et al., 2003; Bheda, 2020). These intrinsic stochastic events are important to consider when populations encounter stressful environments, because phenotypic variation increases the likelihood for survival for some individuals based on the phenotype favoured under certain conditions (Ackermann, 2015; Martins and Locke, 2015).

### Evolutionary advantage of phenotypic heterogeneity

If the different subpopulations originate through stochastic changes in regulatory pathways, then there should be a fitness disadvantage as some members of the population are not ideally suited to the environmental conditions, arguably like the Bottom fraction. However, such variation may have a benefit in environments where the conditions change suddenly, thus one subpopulation unfavoured in one environment is favoured in the new environment, a strategy known as bet-hedging (de Jong et al., 2011; Levy et al., 2012; Grimbergen et al., 2015). Bet-hedging increases a population’s chances of survival, reproductive success, and long-term persistence when environmental conditions are uncertain, fluctuate, or become unfavourable. Without such strategies, isogenic cultures remaining phenotypically identical and respond uniformly to stress could risk extinction if environmental conditions suddenly changed (Martins and Locke, 2015).

In bet-hedging, only a portion of the population varies phenotypically, acknowledging the risks and rewards associated with phenotypic diversity. For example, bacterial persistence, where a subpopulation of persister cells that grow slowly can survive lethal doses of antibiotics (Balaban et al., 2004; Kussell et al., 2005; Smith et al., 2018). Even while growing under steady environmental conditions, a small fraction of persister cells are already present before any antibiotics are applied (Balaban et al., 2004). These persister cells spontaneously switch between normal and slow-growing phenotypes, seemingly to ensure a portion of the population will survive the sudden presence of antibiotics and resume normal growth. This bet-hedging strategy emphasizes how phenotypic diversity helps to spread the risk of the stressor across different phenotypes (Acar et al., 2008; de Jong et al., 2011), and allowing populations to sacrifice optimal fitness in stable conditions for better resilience against environmental stresses (Kussell and Leibler, 2005; Ackermann, 2015).

The phenotypic heterogeneity observed in this study may serve as an adaptive bet-hedging strategy, where subpopulations diversify their phenotypic traits to enhance survival under fluctuating environmental conditions, particularly light. In the case of the Top and Bottom subpopulations, there would be a trade-off between growth and stress resistance. Future work should prioritize an analysis on the expression of genes in these subpopulations to identify expression profiles related to a LL or HL-optimized expression, perhaps like the shared expression patterns linked to fitness trade-offs between growth and stress resistance found in yeast (Kim et al., 2024), supporting the idea that phenotypic diversity arises from stochastic processes that balance competing demands.

## Acknowledgements

This work was supported by a Natural Sciences and Engineering Research Council (NSERC) Discovery Grant (2019-04428) awarded to DGD.

